# Engineering a Synthetic *Escherichia coli* Coculture for Compartmentalized *de novo* Biosynthesis of Isobutyl Butyrate from Mixed Sugars

**DOI:** 10.1101/2023.08.11.553049

**Authors:** Hyeongmin Seo, Gillian Castro, Cong T. Trinh

## Abstract

Short-chain esters are versatile chemicals with use as flavors, fragrances, solvents, and fuels. The *de novo* ester biosynthesis consists of diverging and converging pathway submodules, which is challenging to engineer to achieve optimal metabolic fluxes and selective product synthesis. Compartmentalizing the pathway submodules into specialist cells that facilitate pathway modularization and labor division can present a promising solution. Here, we engineered a synthetic *Escherichia coli* coculture with the compartmentalized sugar utilization and ester biosynthesis pathways to produce isobutyl butyrate from a mixture of glucose and xylose. To compartmentalize the sugar-utilizing pathway submodules, we engineered a xylose-utilizing *E. coli* specialist that selectively consumes xylose over glucose and bypasses the carbon catabolite repression (CCR) while leveraging the native CCR machinery to activate a glucose-utilizing *E. coli* specialist. Upon compartmentalizing the isobutyl butyrate pathway submodules into these sugar-utilizing specialist cells, a robust synthetic coculture could be engineered to selectively produce isobutyl butyrate at a level of 392 mg/L, about 31-fold higher than the monoculture.

## INTRODUCTION

Cellular metabolism comprises of interconnected diverging-converging metabolic pathway modules that are highly regulated^1^. For instance, the fermentable sugar (e.g., glucose or xylose) degradation pathways are converged into the glycolysis to generate pyruvate and then acetyl-CoA that is subsequently diverged to the peripheral metabolic pathways to synthesize cellular resources (e.g., lipids) for cell growth and maintenance^2^. Due to a highly interconnected and regulated metabolic network, introducing a heterologous pathway with diverging-converging pathway modules in a microbial host to achieve optimal product formation has been a challenging but intriguing synthetic biology and metabolic engineering design problem^3^.

One representative diverging-converging biosynthetic pathway is the *de novo* biosynthesis of esters with broad use in many industries such as food, cosmetic, pharmaceutical, and transportation^4^. Microbial biosynthesis of esters uses an alcohol acyltransferase (AAT) that can perform a thermodynamically favorable condensation reaction of an acyl-CoA and an alcohol in an aqueous phase ^5^. The *de novo* ester biosynthetic pathways for converting fermentable sugars to esters can be modularized into submodules, including the sugar-utilizing submodules, acyl-CoA production submodule, alcohol production submodule, and/or ester synthesis submodule^6–8^. Harnessing the modular ester biosynthetic pathways enables combinatorial biosynthesis of a large number of unique esters^6, 7, 9–11^. However, due to the complex nature of diverging-converging ester biosynthetic pathways, it requires optimal metabolic flux balance and control to achieve selective ester biosynthesis^6, 12^.

Isobutyl butyrate is a hydrophobic and sweet-scented medium carbon chain length (C_8_) ester that is naturally found in flowers^13^. A *de novo* isobutyl butyrate biosynthetic pathway is derived from the converging sugar-utilizing pathway submodules to generate pyruvate, then the isobutanol and butyryl-CoA pathway submodules that diverge from pyruvate to make the precursors isobutanol and butyryl-CoA, and finally the ester biosynthesis submodule that converges to synthesize isobutyl butyrate by the AAT-dependent condensation of isobutanol and butyryl-CoA^7^ (Fig. 1A). Although microbial biosynthesis of isobutyl butyrate by an *E. coli* monoculture was previously demonstrated^7, 10^, the production titers were relatively low as compared to acetate ester production such as butyl acetate^12, 14^, isobutyl acetate^15^, isoamyl acetate^9^, and isoprenyl acetate^16^. The common bottlenecks to achieve an efficient isobutyl butyrate production are metabolic burden and optimal expression of the heterologous isobutyl butyrate pathway^6^. Balancing the heterologous isobutanol and butyryl-CoA pathway submodules requires extensive transcription and/or translation optimization of the pathway genes and enzyme solubilization that often perturb cellular resource reallocation such as proteome^9^. One solution to manage a diverging-converging biosynthetic pathway is to deploy a microbial consortium that divides the metabolic labor into two or more specialist cells for various biosynthetic pathways^17^ as demonstrated for production of muconic acid^18^, 4-hydroxybenzoic acid^18^, rosmarinic acid^19^, resveratrol^20^, pyranoanthocyanins^21^, butanol^22^, acetate esters and fatty acid methyl esters (FAME)^23^.

**Figure 1.**
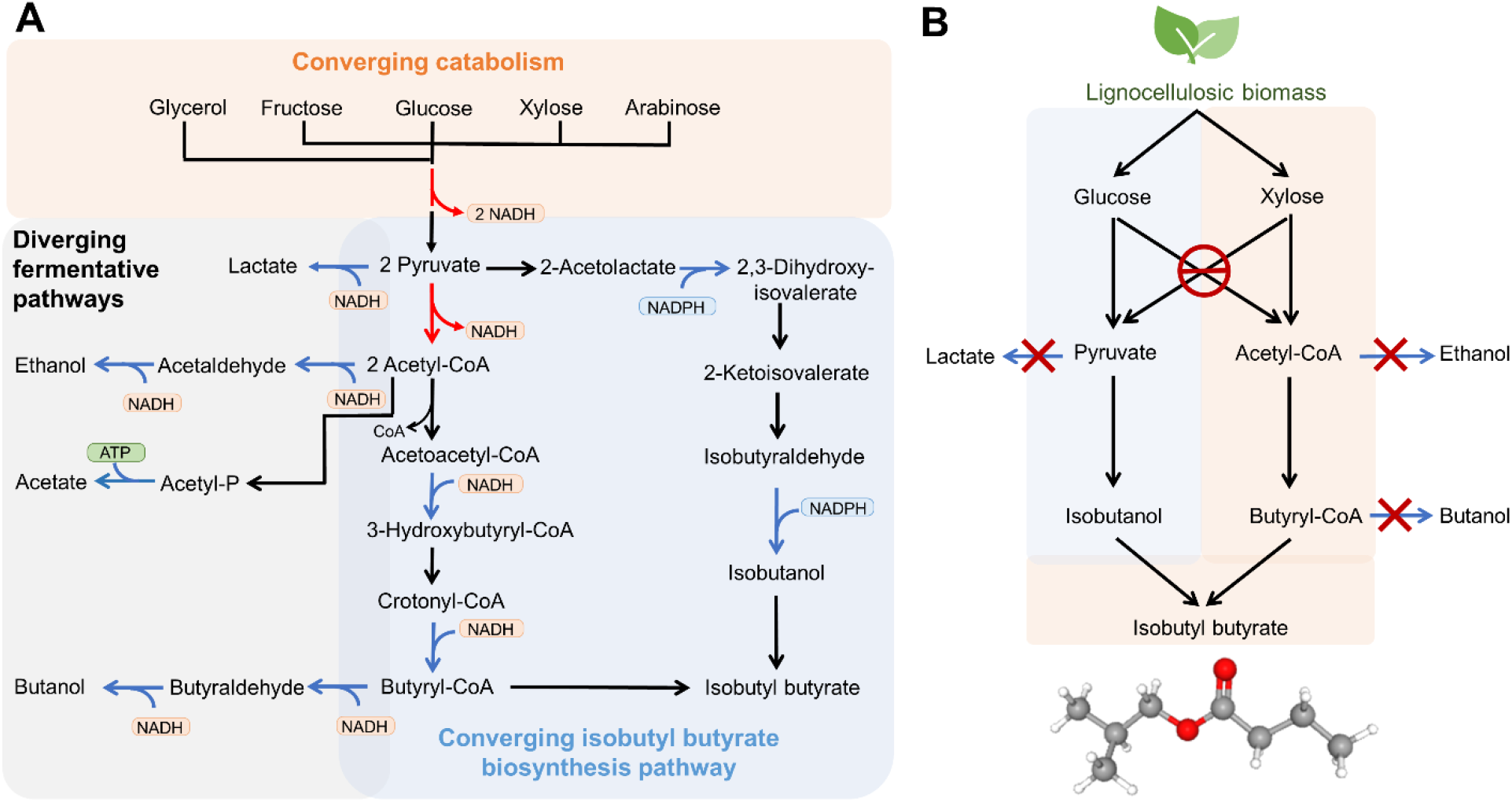
Compartmentalization of the de novo isobutyl butyrate biosynthesis from mixed fermentable sugars. **(A)** The diverging-converging pathways for conversion of mixed sugars to isobutyl butyrate. **(B)** Rational design of compartmentalized sugar utilization and isobutyl butyrate production pathway modules.

In this study, we aimed to engineer and evaluate whether a synthetic coculture of two *E. coli* specialists with the compartmentalized *de novo* isobutyl butyrate biosynthetic pathway overcomes the current ester production limitation in monocultures. We first reprogrammed sugar catabolism of *E. coli* to create sugar-utilizing specialists. Next, we compartmentalized the *de novo* isobutyl butyrate biosynthetic pathway in an engineered *E. coli* coculture of glucose-utilizing and xylose-utilizing specialist cells. We demonstrated the synthetic coculture outperformed the monoculture with improved production and selectivity of isobutyl butyrate.

## RESULTS AND DISCUSSION

### Compartmentalizing sugar catabolism to facilitate simultaneous coutilization of glucose and xylose by an *E. coli* coculture

Glucose and xylose are the most abundant sugars in lignocellulosic biomass, a promising renewable feedstock for biomanufacturing to produce fuels, chemicals, and materials ^24^. While microbial coutilization of glucose and xylose is highly favorable for industrial bioprocessing, most microbes including *E. coli* exhibit a hierarchical sugar preference (i.e., glucose over xylose) due to strong carbon catabolite repression^25^. To bypass this repression, we aimed to compartmentalize the glucose and xylose catabolic pathway submodules to create a coculture of glucose-utilizing and xylose-utilizing *E. coli* specialists that are programmed to convert a mixture of glucose and xylose into isobutyl butyrate (Fig. 1B). To reduce byproduct formation, we employed an *E. coli* BL21(DE3) derived strain HSEC0201^9^, lacking bifunctional alcohol dehydrogenase (*adhE*) and D-lactate dehydrogenase (*dld*) genes, as the initial host for strain engineering (Fig. 1B).

Glucose catabolism of *E. coli* can be manipulated by disrupting the phosphoenolpyruvate sugar transferase system (PTS)^26, 27^ and a glucose transporter (*galP*)^28^. To compartmentalize the glucose and xylose catabolic pathways, we deleted two key glucose catabolic pathway genes, *ptsI* and *glk,* from the chromosome of HSEC0201 to generate HSEC0415 (Fig. 2A). In our design, by leveraging the strong native CCR of *E. coli*, HSEC0201 (or its derivatives without *ptsI* and *glk* deletion) can readily serve as a glucose-utilizing *E. coli* specialist for growth in a mixture of glucose and xylose while HSEC0415 (or its derivatives with *ptsI* and *glk* deletion) functions as a xylose-utilizing *E. coli* specialist. In control experiments, the strain characterization in the glucose-containing M9 medium only showed that the glucose-utilizing parental strain HSEC0201 grew quickly and consumed 9.5 g/L of glucose within 24 hours, as expected (Figs. 2B, 2C). In contrast, the xylose-utilizing strain HSEC0415 cell exhibited significant growth inhibition and only consumed 0.03 g/L of glucose within 50 hours (Figs. 2B, 2C). These results clearly indicated that glucose catabolism of HSEC0415 was drastically inhibited by the *ptsI* and *glk* deletion, as expected and in agreement with previous studies^27, 29^. Although HSEC0415 showed slightly slower cell growth and xylose consumption as compared to HSEC0201, both strains underwent normal growth in the xylose-containing M9 medium (Fig. 2D, E). For growth on a mixture of glucose and xylose, HSEC0201 consumed glucose before consuming xylose (Fig. 2F), exhibiting the hierarchical sugar preference. HSEC0415 showed atypical non-exponential growth within 36 hours (Fig. 2G), which was significantly different from the cell growth in a xylose only medium (Fig. 2D). This result suggested that the glucose-driven CCR clearly limited xylose uptake and inhibited cell growth. Furthermore, HSEC0415 selectively consumed xylose within 36 hours and started to consume glucose, indicating that *ptsI* and *glk* deletions did not eliminate glucose catabolism likely due to other sugar (e.g., mannose) phosphotransferase system^30^ and/or sugar kinases with promiscuous glucose phosphorylation activities^31^.

**Figure 2.**
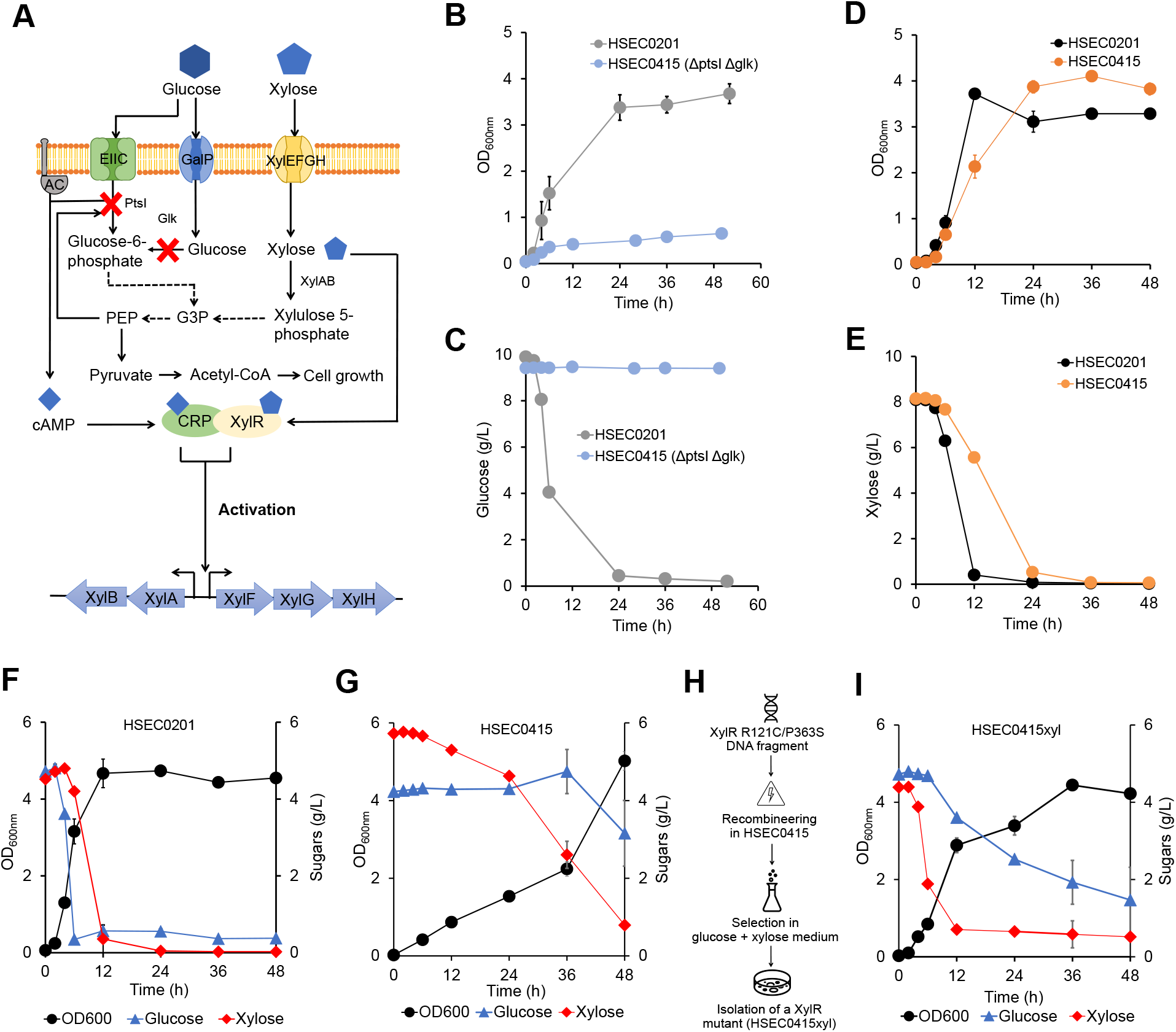
Development of glucose-utilizing and xylose-utilizing *E. coli* specialist to compartmentalize sugar catabolism. **(A)** A scheme of glucose-driven carbon catabolite repression (CCR) against xylose. The red X marks indicate pathway deletion via chromosomal gene deletion. **(B)** Cell growth of the parental strain HSEC0201 and the derived strain HSEC0415 lacking *ptsI* and *glk* genes in the glucose M9 medium. **(C)** Glucose concentration profiles of HSEC0201 and HSEC0415 cell cultures. **(D)** Cell growth of HSEC0201 and HSEC0415 in the xylose M9 medium. **(E)** Xylose concentration profiles of HSEC0201 and HSEC0415 cell cultures. **(F,G)** Kinetic profiles of cell growth, glucose and xylose consumption of **(F)** HSEC0201 and **(G)** HSEC0415. **(H)** Workflow for site-directed chromosomal substitution of *xylR* with *xylR* R121C/P363S. **(I)** Kinetic profiles of cell growth, glucose and xylose consumption of HSEC0415xyl culture in glucose-xylose M9 medium. All the data represent means ± 1 standard deviation from three biological replicates.

To eliminate the glucose-driven CCR against xylose catabolism controlled by the xylose regulator XylR^32^ (Fig. 2A), we substituted the chromosomal *xylR* gene with R121C and P363S mutations that have been shown to confer higher binding affinity to *xyl* operon^33^. We isolated the mutant strain in a glucose-xylose medium through two rounds of liquid culture transfer followed by a dilution plating (Fig. 2H). An isolated strain having chromosomal *xylR* R121C/P363S mutations was designated as HSEC0415xyl. The strain characterization showed that HSEC0415xyl cell growth and xylose uptake were not inhibited by glucose (Fig. 2I), suggesting that *xylR* R121C/P363S resolved the glucose-driven CCR against xylose. Interestingly, HSEC0415xyl culture still remained 0.6 g/L of xylose before consuming glucose. Since HSEC0415xyl completely consumed xylose in the absence of glucose (Fig. 2E), it suggested that glucose might have inhibited transcriptional activation of the *xyl* operon by XylR R121C/P363S at a xylose concentration as low as 0.5 g/L. Overall, by compartmentalizing the sugar catabolism of *E. coli*, we generated glycose-utilizing and xylose-utilizing *E. coli* specialists.

### Rewiring carbon catabolite repression for sugar pathway compartmentalization enabled establishment of a stable coculture of sugar-utilizing *E. coli* specialists

Establishing a robust coculture population is important for efficient biosynthetic pathway compartmentalization and product synthesis. Here, we investigated how the CCR-engineered strains modulated coculture population and heterologous gene expression. To do this, we monitored coculture population dynamics by expressing a green fluorescence protein (GFP) and a red fluorescence protein (RFP) in the engineered *E. coli* cocultures and measuring the fluorescence signals. We introduced an mCHERRY expression plasmid to HSEC0415 (HSEC0415_mCHERRY) and HSEC0415xyl (HSEC0415xyl_mCHERRY) to generate RFP signals. Each of these two strains was paired with a glucose-utilizing *E. coli* specialist partner (i.e., HSEC0302_iLOV) harboring an iLOV expression plasmid that generates GFP signals (Fig. 3A). HSEC0302, derived from HSEC0201, lacks L-lactate dehydrogenase (*ldhA*)^9^, which is beneficial for isobutyl butyrate production pathway compartmentalization with reduced lactate formation. For control experiments, the characterization results in monocultures showed that the iLOV (GFP) and mCHERRY (RFP) fluorescence signals did not interfere with each other (Fig. 3B), suggesting that monitoring the two fluorescence intensities enables dissecting two species in the coculture.

**Figure 3.**
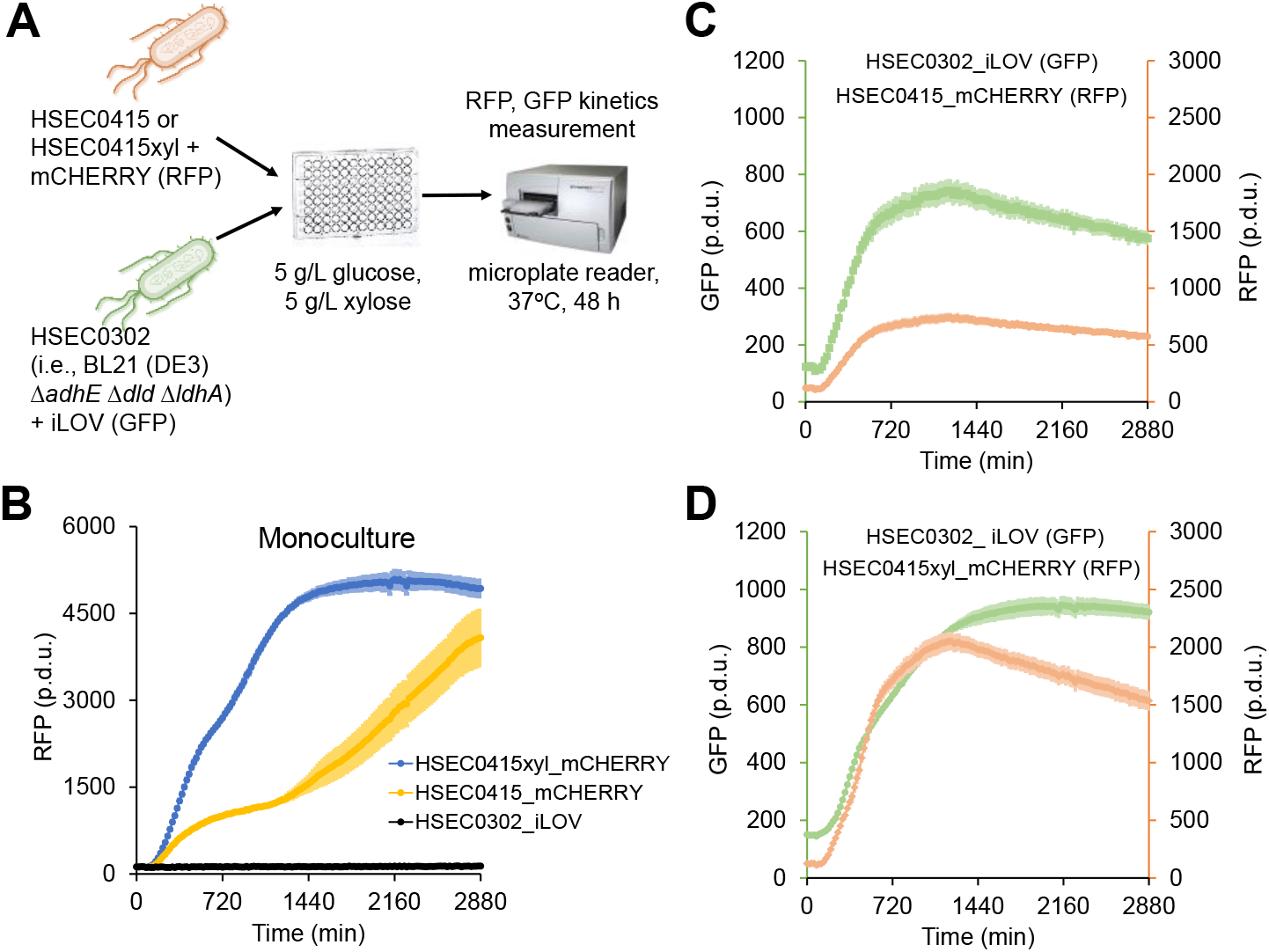
Population dynamics of *E. coli* cocultures with sugar catabolism compartmentalization. **(A)** An experiment scheme of high-throughput cell culture and population dynamics analysis via fluorescence measurement. **(B)** Kinetic profiles of red fluorescence protein (RFP) signals from monocultures. **(C-D)** Green fluorescence protein (GFP) and RFP kinetic profiles of **(C)** HSEC0302_iLOV and HSEC0415_mCHERRY coculture and **(D)** HSEC0302_iLOV and HSEC0415xyl_mCHERRY coculture. All the data represent means ± 1 standard deviation from at least three biological replicates. Abbreviation: p.d.u., procedure defined unit.

We next compared the GFP and RFP kinetics to investigate the compatibility of HSEC0415xyl and HSEC0302 in the coculture (Figs. 3C, 3D). In the control pair of HSEC0415_mCHERRY and HSEC0302_iLOV, the coculture generated the maximum RFP intensity of only 750 procedure defined unit (p.d.u.), which was 5.4-fold lower than the monoculture (4,050 p.d.u.). The RFP signal did not increase after 24 hours as opposed to the monoculture. The results indicated that HSEC0302_iLOV dominated the coculture population in the early phase and inhibited the cell growth of HSEC0415_mCHERRY. In contrast, the RFP and GFP intensities from the coculture of HSEC0415xyl_mCHERRY and HSEC0302_iLOV orthogonally increased at the early phase, stably balancing the population (Fig. 3D). In summary, the sugar pathway compartmentalization enabled generation of glucose-utilizing and xylose-utilizing *E. coli* specialists that form a stable, robust coculture and each can compartmentalize a fluorescent protein production module. The next step is to demonstrate this sugar-utilizing specialist coculture to compartmentalize the diverging-converging isobutyl butyrate production module for enhanced target ester biosynthesis.

### Synthetic Coculture of sugar-utilizing *E. coli* specialists improved isobutyl butyrate biosynthesis

Microbial biosynthesis of isobutyl butyrate in monocultures faces two challenges: i) optimal expression of multiple pathway enzymes with balanced metabolic fluxes for enhanced ester production and ii) control of product selectivity. With the precursors acetyl-CoA, butyl-CoA, ethanol, butanol, and isobutanol available in a monoculture, six possible esters can be formed including: ethyl acetate, butyl acetate, isobutyl acetate, ethyl butyrate, butyl butyrate and isobutyl butyrate due to promiscuity of AATs and AdhE^34–36^. To tackle these challenges, we investigated whether a synthetic coculture of sugar-utilizing *E. coli* specialists with compartmentalized isobutyl butyrate pathway can improve target product synthesis and selectivity. We constructed two *E. coli* strains: the glucose-utilizing *E. coli* specialist HSEC0916 expressing the isobutanol pathway genes and the xylose-utilizing *E. coli* specialist (HSEC1017 expressing the butyryl-CoA pathway genes (Fig. 4A). Since the native AdhE gene was deleted in the engineered sugar-utilizing *E. coli* specialists, the formation of ethyl esters (ethyl acetate, ethyl butyrate) and butyl esters (butyl acetate and butyl butyrate) can be avoided. Because butyryl-CoA is an intracellular metabolite, we expressed AATs in HSEC1017 to condense butyryl-CoA into isobutyl butyrate (Fig. 4B). We deployed two butyrate ester forming AATs, including SAAT^35^ and CATec3 Y20F^36^, to accelerate isobutyl butyrate production (Fig. S1A). Due to the substrate promiscuity of AATs^36, 37^, isobutyl acetate byproduct is inevitable in our design.

**Figure 4.**
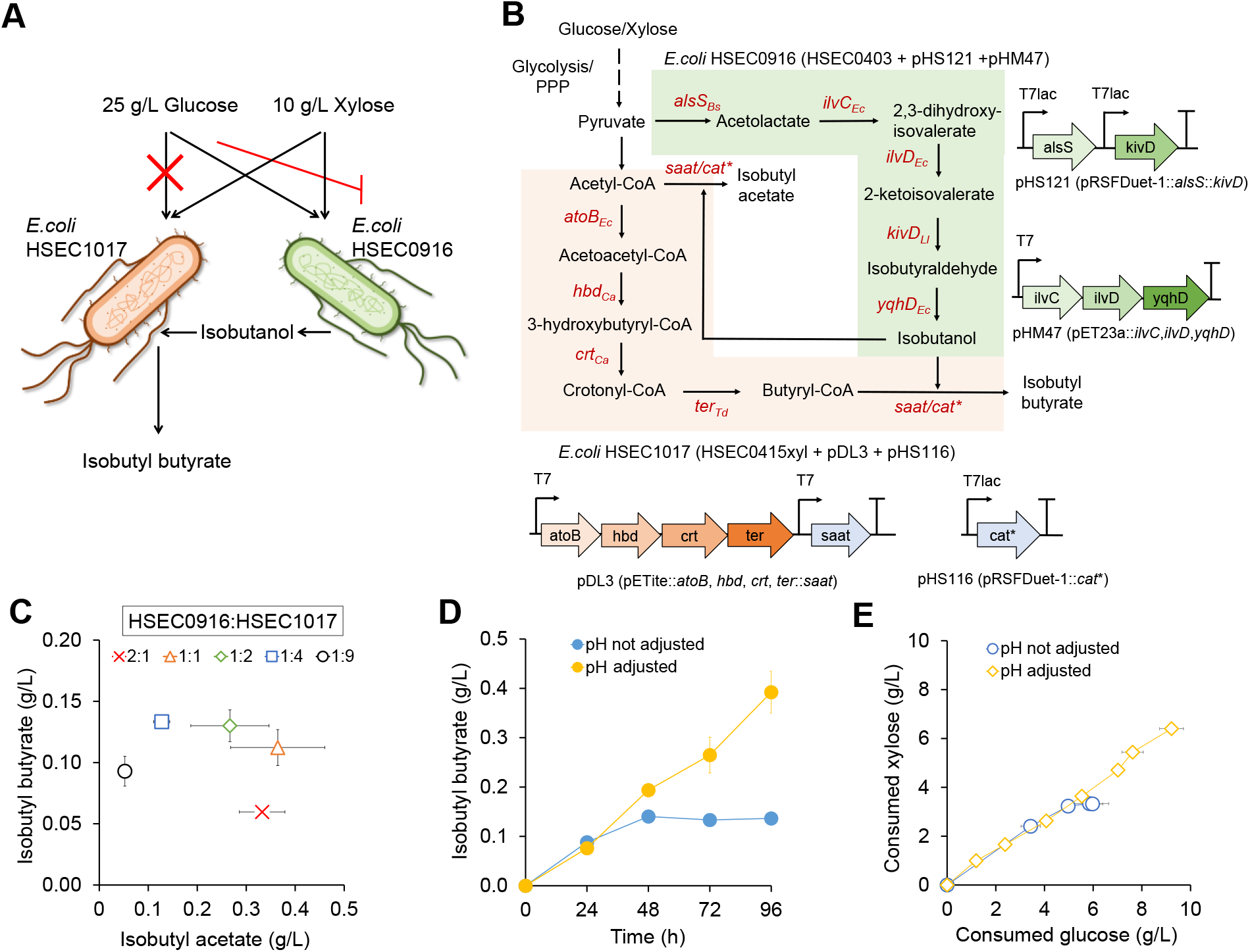
Compartmentalization of the *de novo* isobutyl butyrate production pathway by a synthetic coculture of glucose-utilizing and xylose-utilizing *E. coli* specialists. **(A)** Design of the *de novo* isobutyl butyrate microbial biosynthesis from a mixture of glucose and xylose using a synthetic *E. coli* coculture. **(B)** Designs of metabolic pathway and synthetic operons for isobutyl butyrate production from mixed fermentable sugars. Abbreviations: Bs, *Bacillus substilis*; Ec, *Escherichia coli*; Ll, *Lactococcus lactis;* Ca, *Clostridium acetobutylicum;* Td, *Treponema denticola;* PPP, pentose phosphate pathway. **(C)** Titers of isobutyl butyrate and isobutyl acetate from cocultures with various inoculation ratios. **(D)** Kinetic profiles of isobutyl butyrate production with and without pH adjustment. **(E)** Sugar consumption by a coculture of HSEC0916 and HSEC1017 with and without pH adjustment. All the data represent means ± 1 S.D. from three biological replicates.

We started characterizing the production of isobutyl butyrate from the coculture of HSEC1017 and HSEC0916 at various HSEC1017: HSEC0916 inoculum ratios of 2:1, 1:1, 1:2, 1:4, and 1:9 in a glucose-xylose medium (Fig. 4C). To simulate a sugar mixture from lignocellulose biomass hydrolysates^38^, 25 g/L glucose and 10 g/L xylose were added as carbon and energy sources. Our results showed that the 1:1 inoculum ratio achieved the highest total ester production of 476 mg/L including isobutyl acetate and isobutyl butyrate; however, the selectivity of isobutyl butyrate with respect to isobutyl acetate was as low as 13% (mol/mol). The higher HSEC0916 inoculation ratio produced more isobutyl acetate, while isobutyl butyrate titers were not significantly changed. The inoculum ratio of 1:4 achieved the highest isobutyl butyrate selectivity of 46% (mol/mol) with the highest titer of 133 ± 14 mg/L (Fig. 4C), which were 10.5-fold higher than *E. coli* monoculture reported previously (12.6 mg/L)^7^. The significantly improved titer and product selectivity suggested that compartmentalizing the isobutanol and butyryl-CoA pathways into the two strains was beneficial for isobutyl butyrate production. The two byproducts isobutanol and isobutyl acetate were produced at the titers of 521 ± 25 mg/L and 127 ± 15 mg/L, respectively, indicating that butyryl-CoA and/or AAT pathway submodules were the bottlenecks of the isobutyl butyrate production.

During the strain characterization, we observed that the pH rapidly decreased below 5 after 24 hours, impeding continuation of isobutyl butyrate production (Fig. S1B). Under the pH maintained between 6.0 and 7.0, isobutyl butyrate production titers reached up to 392 mg/L after 96 hours (Fig. 4D). Especially, the coculture maintained steady isobutyl butyrate production and sugar consumption with linear glucose and xylose consumption (Fig. 4E). A linear regression of the consumption ratio exhibited 0.68 (g xylose /g glucose) with a high coefficient of determination (R^2^=0.99), further supporting the conclusion that the coculture of HSEC1017 and HSEC0916 stably maintained the population with orthogonal glucose and xylose consumption.

Overall, the *de novo* isobutyl butyrate biosynthetic pathway was successfully compartmentalized by a synthetic coculture, enabling simultaneous conversion of glucose and xylose into isobutyl butyrate. Production of isobutyl butyrate can be modulated by controlling pH and inoculation of cocultures with the compartmentalized isobutyl butyrate pathway module.

## CONCLUSIONS

Synthetic microbial consortia can perform complex tasks that are difficult or impossible for individual species, such as simultaneous consumption of mixed sugars, efficient expression and flux balance of diverging-converging pathways, and control of product selectivity^39^. In this study, we rationally designed sugar-utilizing *E. coli* specialist cocultures to compartmentalize both the diverging-converging sugar-utilizing and isobutyl butyrate producing pathways for efficient conversion of mixed sugars into isobutyl butyrate. The improved isobutyl butyrate titer and selectivity from glucose-xylose co-utilization suggests that a synthetic microbial consortium can serve as a novel biomanufacturing platform for ester overproduction. Recent discovery of CATs that can be repurposed to function as efficient and robust AATs^36^ makes it highly feasible to create synthetic microbial consortium to synthesize esters. Since microbial consortia are capable of assimilating various renewable feedstocks, we envision the ester pathway compartmentalization strategy can be widely applicable to develop microbial consortia for effective production of designer esters.

## METHODS

### Media and cultivation

*E. coli* strains were grown in lysogeny broth (LB) medium or a modified M9 medium containing 5 g/L yeast extract and 10 g/L MOPS. Either glucose or a mixture of glucose and xylose were included in the M9 medium as specified elsewhere. Antibiotics were supplemented with 100 µg/mL ampicillin and/or 50 µg/mL kanamycin when appropriate. For heterologous gene expression and ester production, 0.1 mM of isopropyl β-d-1-thiogalactopyranoside (IPTG) was added to the media before inoculating cells. The sugar catabolism characterization experiments were performed in a 125 mL flask with a 10 mL working volume of M9 medium. For the 96-well microplate-based cell culture, cells were cultured in M9 medium with a150 μL working volume. For coculture population analysis, RFP and GFP signals were monitored at Ex_530nm_/Em_590nm_ and Ex_485nm_/Em_528nm_, respectively.

For the isobutyl butyrate production, cells were cultured in a 125 mL screw-capped shake flask with a 20 mL working volume of M9 medium containing 25 g/L glucose and 10 g/L xylose. 10 mL of hexadecane was overlaid to extract ester products during the culture. For the pH adjusted batch culture, 50 μL-100 μL of 5M KOH was intermittently added every 12 h to maintain pH between 6.0 and 7.5.

### Strain and plasmid construction

#### Plasmid construction

The list of strains and plasmids used in this study is presented in Table 1. Plasmids were constructed by Gibson DNA assembly using the primers listed in Table S1. Phusion DNA polymerase (cat# F530S, Thermo Fisher Scientific, MA, USA) and DNA purification and gel extraction kits (Omega Biotek, GA, USA) were used to amplify DNA fragments and purify the PCR products. *E. coli* TOP10 was heat-shock transformed after being mixed with the assembled DNA and selected on LB agar plates containing appropriate antibiotics. All the constructed plasmids were confirmed by Sanger sequencing.

**Table 1:** List of plasmids and strains used in this study.

| Name | Descriptions | Source |
| --- | --- | --- |
| **Plasmids** |  |  |
| pET23a | pBR322 ori, Amp<sup>R</sup>, lacI, T7lac promoter | Novagen |
| pETite* | pBR322 ori, Amp<sup>R</sup>, lacI, T7lac promoter | Novagen |
| pRSFDuet-1 | RSF ori, Kan<sup>R</sup>, lacI, T7lac promoter | Novagen |
| pSIM6 | pSC101 repA<sup>ts</sup>, Gam, Beta, Exo under the control of a temperature sensitive promoter | 41 |
| pCP20 | repA101<sup>ts</sup>, Cm<sup>R</sup>, Amp<sup>R</sup>, FLP recombinase under the control of a temperature sensitive promoter. | 43 |
| pET_iLOV | pET29a::iLOV (GFP) | This study |
| pET_mCHERRY | pET29a::mCHERRY (RFP) | This study |
| pHM47 | pET23a::ilvC, ilvD, yqhD | 44 |
| pHS121 | pRSFDuet-1::alsS::kivD | 9 |
| pDL3 | pETite*::atoB, hbd, crt, ter::saat | 7 |
| pHS116 | pRSFDuet-1::catec3 Y20F | This study |
| **E. coli strains** |  |  |
| Top10 | Host for molecular cloning, mcrA, Δ(mrr-hsdRMS-mcrBC), Phi80lacZ(del)M15, ΔlacX74, deoR, recA1, araD139, Δ(ara-leu)7697, galU, galK, rpsL(SmR), endA1, nupG | Invitrogen |
| BL21 (DE3) | Host for plasmid in vivo methylation, E. coli B dcm, ompT, hsdS(rB-mB-), gal | Invitrogen |
| HSEC0201 | BL21(DE3) ΔadhE, Δdld | 9 |
| HSEC0302 | BL21(DE3) ΔadhE, Δdld, ΔldhA | 9 |
| HSEC0415 | BL21(DE3) ΔadhE, Δdld, ΔptsI, Δglk | This study |
| HSEC0415xyl | BL21(DE3) ΔadhE, Δdld, ΔptsI, Δglk, xylR::xylR R121C/P363S | This study |
| HSEC0916 | HSEC0403 harboring pHS121 and pHM47 | This study |
| HSEC1017 | HSEC0415xyl harboring pHS116 and pDL3 | This study |

#### Recombineering

*E. coli* gene deletions and XylR site-directed substitution were carried out via recombineering by utilizing a temperature-sensitive recombineering toolkits^40, 41^, as described previously^9^. For HSEC0415xyl construction, antibiotics-free selection was performed. After electroporating the linear fragment of *xylR* R121C/P363S, the cells were recovered in a liquid LB medium at 30℃ for 2 hours. Then, the cells were inoculated in 10 mL M9 hybrid medium containing 10 g/L glucose and 10 g/L xylose at 1% (v/v) inoculum ratio and cultured overnight. 100 μL of the cells was passaged to a fresh 10 mL M9 hybrid medium containing 10 g/L glucose and 10 g/L xylose, followed by overnight shaking incubation at 37°C and 200 rpm. The grown cells were serial diluted from 10^-6^ to 10^-8^ and plated on a LB medium containing 10 g/L glucose and 10 g/L xylose overnight. The *xylR* region of eight colonies was PCR amplified followed by Sanger sequencing. Five out of the eight colonies (62.5%) showed the correct substitution of *xylR* with *xylR* R121C/P363S.

### Analytical methods

#### High-performance liquid chromatography (HPLC) analysis

HPLC system (Shimadzu Inc., MD, USA) was used to quantify extracellular metabolites and sugars. For HPLC sample preparation, 800 μL of culture samples were centrifuged at 17,000 xg for 3 minutes followed by filtering through 0.2 micron filters. The samples were run with 5 mM H_2_SO_4_ at 0.6 mL/min on an Aminex HPX-87H column (Biorad Inc., CA, USA) at 50℃. A dual refractive index detector (RID) and ultra-violet detector (UVD) at 220 nm were used to identify and determine analyte concentrations.

#### Gas chromatography coupled with mass spectroscopy (GC/MS) analysis

GC (HP 6890, Agilent, CA, USA) equipped with a MS (HP 5973, Agilent, CA, USA) was used to quantify isobutyl acetate and isobutyl butyrate as described previously^36^. A Zebron ZB-5 capillary column (Phenomenex, CA, USA) was used with He as the carrier gas at a flow rate of 0.5 mL/min. The oven temperatures were programed as 50°C initial temperature, 1°C/min ramp up to 58°C, 25°C/min ramp up to 235°C, 50°C/min ramp up to 300°C, and 2-minutes bake-out at 300°C. 1 μL hexadecane layer sample was injected into the column with the splitless mode at an injector temperature of 280°C. For the MS system, selected ion mode (SIM) was used to detect and quantify target esters with the parameters described previously^42^. For every analysis batch, external and internal standards were used to quantify the analytes. As an internal standard, 10 mg/L n-decane was added in initial hexadecane layer and detected with m/z 85, 99, and 113 from 12 to 15 minute retention time range.

## ACKNOWLEDGEMENTS

This research was funded by the DOE BER award (DE-SC0022226), the DOE subcontract grant (DE-AC05-000R22725) from the Center of Bioenergy Innovation (the DOE Bioenergy Research Center funded by the Office of Biological and Environmental Research in the DOE Office of Science), and the DOE Joint Genome Institute. The work conducted by the U.S. Department of Energy Joint Genome Institute, a DOE Office of Science User Facility, is supported under Contract No. DE-AC02-05CH11231. The authors would like to thank the Center of Environmental Biotechnology at UTK for using the GC/MS instrument.

## AUTHOR CONTRIBUTIONS

Conceptualization: CT; Data Curation: HS; Formal Analysis: HS; Funding Acquisition: CT; Investigation: HS, GC; Methodology: HS; Project Administration: CT; Resources: CT; Supervision: CT; Validation: HS, GC; Visualization: HS, CT; Writing: HS; Writing-Review & Editing: HS, CT.

## SUPPLEMENTARY MATERIALS

**Supplementary File** contains Table S1 and Figures S1.

**Table S1.** A list of primers used in this study. The bold and underlined letters indicate restriction enzyme recognition sites and site-directed DNA sequence changes, respectively.

Figure S1. Optimizing isobutyl butyrate production by the coculture of HSEC0916 and HSEC1017 at inoculum ratio of 1:4. **(A)** Final isobutyl butyrate titers by overexpression of CATec3 Y20F and co-expression of CATec3 Y20F and SAAT. **(B)** A profile of isobutyl esters without pH adjustment. Each data represents means ± 1 standard deviation from three biological replicates.

## Notes

### Competing Interest Statement

The authors have declared no competing interest.

