## Supplementary Materials for "Engineering a Synthetic *Escherichia coli* Coculture for Compartmentalized *de novo* Biosynthesis of Isobutyl Butyrate from Mixed Sugars"

**Table S1.** A list of primers used in this study. The bold and underlined letters indicate restriction enzyme recognition sites and site-directed DNA sequence changes, respectively.

| Primer name | Primer sequence (5’ to 3’) | Description |
|---|---|---|
| HS847 | GGTTCTTTTAAAAATCAGTCACAAGTAAGGTAGGGTTATGGTGTAGGCTGGAGCTGCTTC | *ptsI* deletion forward |
| HS848 | TGATCTTCTCCTAAGCAGTAAATTGGGCCGCATCTCGTGGACATATGAATATCCTCCTTA | *ptsI* deletion reverse |
| HS849 | AAGACGAGCAGAAAGCGG | *ptsI* check upstream |
| HS850 | AGTTCCGGTATCCTTCTTG | *ptsI* check downstream |
| HS843 | GAAAGAATTATTTTGACTTTAGCGGAGCAGTTGAAGAATGGTGTAGGCTGGAGCTGCTTC | *glk* deletion forward |
| HS844 | GATTATCGGGGAGAGTTACCTCCCGATATAAAAGGAAGGATCATATGAATATCCTCCTTA | *glk* deletion reverse |
| HS845 | GATATTTACAGGGAGCCTGCCTTTC | *glk* check upstream |
| HS846 | ACGCAGGTCGGCCTTGTG | *glk* check downstream |
| HS740 | TAACCTAGGCTGCTGCCACCGCTGAGCAAT | pRSFDuet-1 backbone for pHS116 forward |
| HS739 | GTGGTGATGATGGTGATGGCTGCTGCCCATGG | pRSFDuet-1 backbone for pHS116 reverse |
| HS741 | CCATGGGCAGCAGCCATCACCATCATCACCAC<br>ATGAATTATACTAAATTCGATG | CATec3 Y20F for pHS116 forward |
| HS742 | ATTGCTCAGCGGTGGCAGCAGCCTAGGTTA<br>TTACTTCAATTTCGAATTGCAG | CATec3 Y20F for pHS116 reverse |
| HS902 | CTCT**GAGCTC**AGGAAAAGAACCATGTTTACTAAACGTCAC | XylR cloning in a plasmid forward |
| HS903 | AACCA**GCGGCCGC**CTACAACATGACCTCGCTATTTACATC | XylR cloning in a plasmid reverse |
| HS904 | CGTTAACTGTTTTGCTTTTTATGGTCTTCCG | XylR R121C mutagenesis forward |
| HS905 | AAGCAAAACAGTTAACGCCTTTCTCTTTTAAATGC | XylR R121C mutagenesis reverse |
| HS906 | CGGTTATTCCTCGCTGCAATATTTCTACTCTG | XylR P363S mutagenesis forward |
| HS907 | GCAGCGAGGAATAACCGCACATTTGCGATATC | XylR P363S mutagenesis reverse |
| HS908 | AGGAAAAGAACCATGTTTACTAAACGTCAC | XylR PCR for recombineering forward |
| HS909 | CTACAACATGACCTCGCTATTTACATC | XylR PCR for recombineering reverse |

**Figure S1.** Optimizing isobutyl butyrate production by the coculture of HSEC0916 and HSEC1017 with an inoculum ratio of 1:4. **(A)** Final isobutyl butyrate titers by overexpression of CATec3 Y20F and co-expression of CATec3 Y20F and SAAT. **(B)** A kinetic profile of isobutyl esters without pH adjustment. Each data represents means ± 1 standard deviation from three biological replicates.

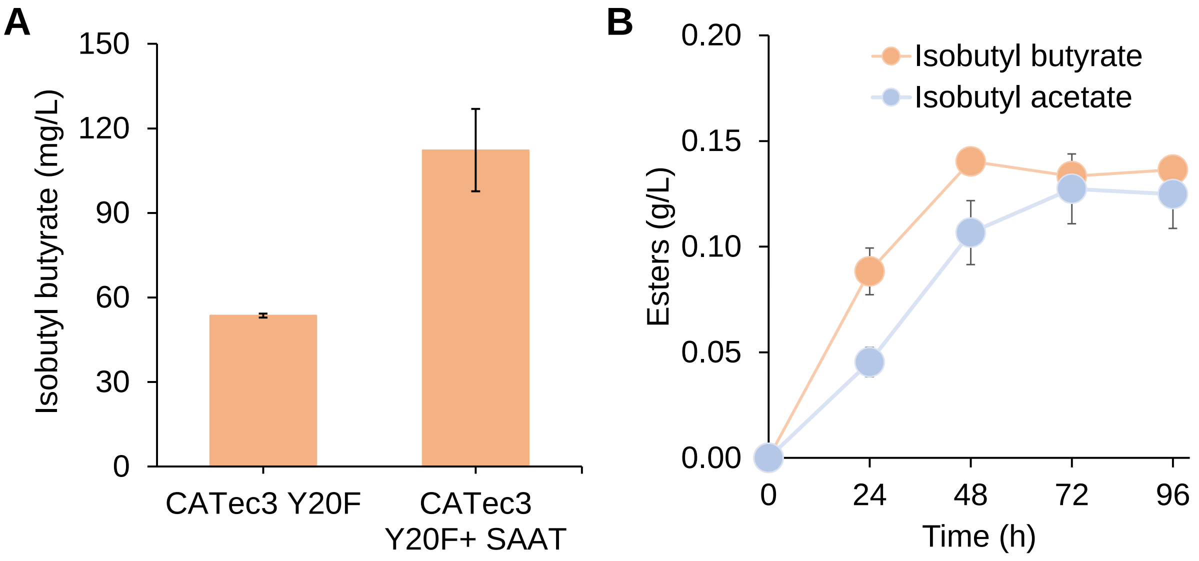
